# Loss of WNT2B Increases Progression from Dysplasia to Colorectal Cancer

**DOI:** 10.1101/2025.08.21.671161

**Authors:** Luiz Fernando Silva Oliveira, Yu-Syuan Wu, Sathuwarman Raveenthiraraj, Jaedeok Kwon, Venkata Siva Dasuri, Comfort Adegboye, Juan Putra, Jorge O. Munera, Diana L. Carlone, David T. Breault, Amy E. O’Connell

**Author notes:** **Corresponding Authors:** Luiz Fernando Silva Oliveira, Ph.D., Address: Enders 9, 300 Longwood Ave, Boston, MA 02115.; Amy E. O’Connell, MD, Ph.D., Address: Enders 9, 300 Longwood Ave, Boston, MA 02115. **Support**: This work was supported by the National Institutes of Health NIDDK K08DK120871 (A.E.O.), NIH P30DK034854-36 (Harvard Digestive Disease Center), and the Division of Newborn Medicine at Boston Children’s Hospital. **Conflict of interest:** The authors declare no conflicts of interest.

## Abstract

Colorectal cancer (CRC) is the third most common cancer and the second leading cause of cancer-related deaths in the United States, and upregulation of the WNT pathway is a primary driver in most cases. However, the role of individual WNT proteins in the development of CRC remains poorly understood. Our previous studies demonstrated that WNT2B loss-of-function leads to severe intestinal enteropathy in humans and increases chemically-induced colitis in mice, suggesting a protective function in the colon. Therefore, we investigated how loss of WNT2B affects CRC development. We used azoxymethane (AOM)/dextran sodium sulfate (DSS) to model colitis-associated cancer (CAC) and AOM-induced mutagenesis to model sporadic CRC. We measured the number and size of tumors and performed histopathological and molecular analyses. We also analyzed the Cancer Genome Atlas to evaluate *WNT2B* expression in human colon cancer. In CAC and CRC mouse models, *Wnt2b KO* mice showed decreased survival and enhanced tumor burden. Moreover, *Wnt2b KO* mice had larger tumors and enhanced dysplasia, with a higher frequency of animals progressing from adenomas to adenocarcinomas compared to control littermates. *Wnt2b KO* animals frequently presented with intestinal bleeding and rectum prolapse, which resembles obstructive CRC. Furthermore, *WNT2B* expression was downregulated in human CRC samples compared to healthy controls, which predicted a significantly lower patient survival. These findings support the conclusion that WNT2B is required for maximal resistance against tumorigenesis and raise the possibility that selectively increasing WNT2B signaling may be a useful colon cancer prevention strategy.

**Significance:** WNT2B loss-of-function increases colon cancer tumorigenesis. Targeting WNT2B may represent a novel strategy for intestinal diseases with a high risk of neoplastic transformation, potentially decreasing the progression to cancer development.

## Introduction

Colorectal cancer (CRC) is the third most common cancer diagnosed and the second leading cause of cancer-related deaths in both men and women in the United States (1, 2). Hyperactivation of the Wingless-related integration site protein (WNT) pathways is the driver of nearly all cases of CRC (3). However, the role of individual WNT proteins in the development of CRC remains poorly understood. WNT signals via canonical (β-catenin-dependent) and non-canonical (β-catenin-independent) pathways to regulate the expression of genes involved in different aspects of cell behavior, such as proliferation, differentiation, and stemness (4). Nineteen different secreted proteins known as WNT ligands have been identified in humans, including WNT2B (5).

We previously demonstrated that WNT2B loss-of-function (LOF) is associated with abnormal intestinal morphology in humans and decreased expression of intestinal stem cell markers in humans and mice (6–8). Clinically, WNT2B LOF causes severe congenital enteropathy in humans (6), while *Wnt2b KO* mice have enhanced susceptibility to early injury in a mouse model of acute colitis due to increased immune cell recruitment and proinflammatory cytokine production (8). Another group has recently reported that human WNT2B LOF also predisposes to gastrointestinal dysplasia as affected subjects age (9).

Chronic inflammation is a cancer hallmark, and patients with inflammatory bowel disease (IBD) are at increased risk of developing a subtype of CRC known as colitis-associated cancer (CAC) (10). In contrast, sporadic CRC features overactivation of the canonical WNT pathway due to alterations in intracellular regulators like adenomatous polyposis coli (APC), AXIN, and β-catenin (3). Therefore, targeting upstream WNT effectors has been considered irrelevant or expected to have no benefit in CRC (5). However, these molecular alterations are described as late-onset events in other CRC subtypes, such as CAC, revealing a heterogeneous WNT signaling function during the different stages of tumor development. Nonetheless, other groups have shown that secreted extracellular agonists (i.e., WNTs and R-spondins) and antagonists (i.e., NOTUM, WIFI, and DKK) are involved in CRC progression and are associated with more aggressive tumor phenotypes (5). Still, the role of WNT2B in the context of CRC development has not been characterized, and recent work suggests that WNT2B may not be a canonical Wnt, as previously thought (11). Here we examine the effects of WNT2B LOF in the context of CRC using mouse models of inflammation-associated tumorigenesis and sporadic tumorigenesis via a mutagenesis-induced cancer model.

## Materials and Methods

### Research Ethics

All animal experiments were approved by Boston Children’s Hospital’s Institutional Animal Care and Use Committee (IACUC) under protocol #00001978.

### Rigor and Reproducibility

Experimental data from mouse experiments represent a minimum of six biological replicates, which, based on our previous calculations using an alpha of 0.05, gives adequate power to detect meaningful differences between control littermates and *Wnt2b KO* mice.

### Reagents

Supplementary Table 1 describes all reagents used in this study, including manufacturer, catalog numbers, antibody clones, and concentrations.

### Mice

*Wnt2b KO* mice and control littermates were generated as previously reported, with no significant differences found when comparing *Wnt2b* expression or experimental outcomes between wild-type (WT) and heterozygous littermates (8). Therefore, *Wnt2b* WT and heterozygous individuals were combined into the control group throughout the manuscript. All mice were maintained in micro-isolator cages with free access to food and water under a 12-hour light/12-hour dark cycle at the Animal Resources at Boston Children’s Hospital (ARCH).

### Colitis-Associated Cancer Model

We adapted the well-established AOM/DSS model of CAC, given our previous study showing that *Wnt2b KO* mice have increased sensitivity to DSS (8) and would likely die from the treatment if we followed the standard 5–7-day colitis cycle (12, 13). However, we included an extra DSS cycle to facilitate tumor development. On experimental day 1, male and female control and *Wnt2b KO* mice aged 8-12 weeks received a single intraperitoneal (i.p.) injection of AOM (Sigma-Aldrich) (dose = 10 mg/kg of body weight). Five days later, we treated the mice with 2.5% DSS (MPBio) in their drinking water for three days. The mice were then allowed to recover for sixteen days, and this regimen was repeated four times. Weight loss, stool consistency, and rectal bleeding were scored daily to generate a semiquantitative clinical colitis assessment (12, 13). See Supplemental Table 2 for detailed Disease Activity Index (DAI) scores. Mice were euthanized on day 75 or if they met criteria for early euthanasia, which included extreme distress, hunching, rectal prolapse, or if they lost more than 20% of their initial body weight.

### Sporadic Colorectal Cancer Model

Male and female *Wnt2b* control and *Wnt2b KO* mice aged 8-12 weeks were treated weekly with an i.p. injection of 10 mg/kg AOM for six consecutive weeks to mimic sporadic colorectal cancer (14, 15). Changes in body weight, feces consistency, rectal bleeding, or any signs of distress or behavioral changes were recorded daily. Mice were euthanized 12 weeks after the first AOM injection or sooner whenever they presented extreme distress and met criteria for early euthanasia, which included hunching, rectal prolapse, or losing more than 20% of their initial body weight.

### Histology

After euthanasia, colons were removed, measured, cleaned, and opened longitudinally. The tissue was then Swiss-rolled, placed in fixation cassettes, and fixed for 24 hours in 10% Neutral-Buffered Formalin at room temperature (RT). Following routine histology processing, the formalin-fixed paraffin-embedded (FFPE) tissue samples were sectioned at 5 µm depth for H&E or PAS staining. Histologic assessment for colitis, dysplasia, and tumor staging was done by a pathologist (J.P.) who was blinded to the study design. H&E- and PAS-stained slides were evaluated for mucosal and submucosal inflammatory cell infiltrates, epithelial abnormalities (crypt hyperplasia and goblet cell loss), crypt loss, ulceration, and crypt abscess. Dysplasia was classified as low-grade or high-grade, depending on nuclear atypia (polarization, stratification, and chromatism), epithelial differentiation, and abnormal growth patterns (16, 17). Detailed histopathological scores are described in Supplemental Table 3.

### Digital Image Analysis

Whole-slide image analysis of H&E-stained Swiss rolls from mouse colons was performed using QuPath (version 0.5.1) (18). Tile scan images were obtained at 20X magnification using an EVOS FL Auto Imaging System (Thermo Fisher Scientific). Images were opened, and tumor regions previously identified in the histopathological analysis were manually annotated using the freehand selection method, as previously described (19). Annotations were stored as region-of-interest (ROI) objects. Using established criteria, tumors were then classified by anatomical location (proximal, middle, and distal) and stage (low-grade and high-grade adenoma, or adenocarcinoma) (16, 17). Tumor area (µm²) was calculated within QuPath, and measurement data were exported. Area values were converted to mm² in Microsoft Excel using a custom macro (18, 19).

### RNA Purification and Real-Time Polymerase Chain Reaction

Distal colon samples (1 cm) were obtained from untreated controls and AOM/DSS-treated animals. Samples were digested with TRIzol™ Reagent (Invitrogen), and RNA was purified using the RNeasy Mini Kit (Qiagen), according to the manufacturer’s instructions. RNA was quantified with a NanoDrop (Invitrogen) and reverse transcribed into cDNA using a high-capacity cDNA reverse transcription kit (Thermo-Fisher) per the manufacturer’s instructions. Gene expression was analyzed via real-time polymerase chain reaction, performed on a QuantStudio6 Flex (Thermo-Fisher) using the TaqMan qPCR Master mix and specific primers as specified in Supplemental Table 4. Technical duplicates or triplicates were implemented to account for technical variability, with 18S as an internal control, and fold change was calculated using the ΔΔCt method.

### Bioinformatic and Statistical Analyses

Human colorectal cancer genomic and transcriptomic data from The Cancer Genome Atlas (TCGA) were analyzed using online bioinformatic platforms. WNT2B gene expression profiles from healthy and colorectal cancer samples were analyzed using the TNMplot (20) and Gene Expression Profiling Interactive Analysis (GEPIA) (21). KM Plotter (22) was used to determine the association between WNT2B expression levels and patient survival. Genomic analysis was performed using cBioPortal (23). Statistical analyses were performed using GraphPad Prism 10 or R software. All data were tested for normality using the Shapiro-Wilk test. A P value of less than 0.05 was considered statistically significant for all downstream analyses. Tumor incidence and sizes between Control and *Wnt2b KO* mice were compared by two-sided Fisher’s exact test and unpaired two-tailed t-test, respectively. Gene expression was compared using two-way ANOVA with Tukey’s post hoc test. Kaplan-Meier survival analysis was used to predict the significance of WNT2B expression in CRC patients and mouse experiments, with significance determined using the log-rank test.

### Data Availability Statement

The data generated in this study are available within the article and its supplementary data files, and the raw data are available by email to the corresponding author.

## Results

### *Wnt2b KO* Mice Are More Susceptible to AOM/DSS-induced CAC

Using an adapted model of AOM/DSS (**Fig. 1A**), we observed a significant decrease in body weight in *Wnt2b KO* mice during the first and last DSS cycles (**Fig. 1B**). Despite only three days of DSS treatment, several *Wnt2b KO* male mice reached maximum body weight loss above 20% (**Fig. S1A**), which resulted in significantly decreased survival compared to control littermates (**Fig. 1C**). We also observed progressive intestinal bleeding (**Fig. 1D**). Consequently, the disease activity index (DAI), which is based on weight loss, stool consistency, and intestinal bleeding (12, 13) was significantly higher for *Wnt2b KO* mice compared to controls throughout the experiment (**Fig. S1B**). At the end of the experimental protocol, no significant differences were observed in colon length (**Fig. S1C**) or spleen weight (**Fig. S1D**). On the other hand, one or more macroscopic tumors were observed in 100% (9/9) of *Wnt2b KO* mice surviving the whole protocol compared to ∼75% (14/18) of control animals (**Fig. 1E**). Furthermore, *Wnt2b KO* mice had a significantly enhanced tumor burden, or number of tumors per mouse, than control littermates (**Fig. 1F**), with scattered neoplastic lesions predominantly identified in the distal colon (**Fig. 1G-H**).

**Fig. 1.**
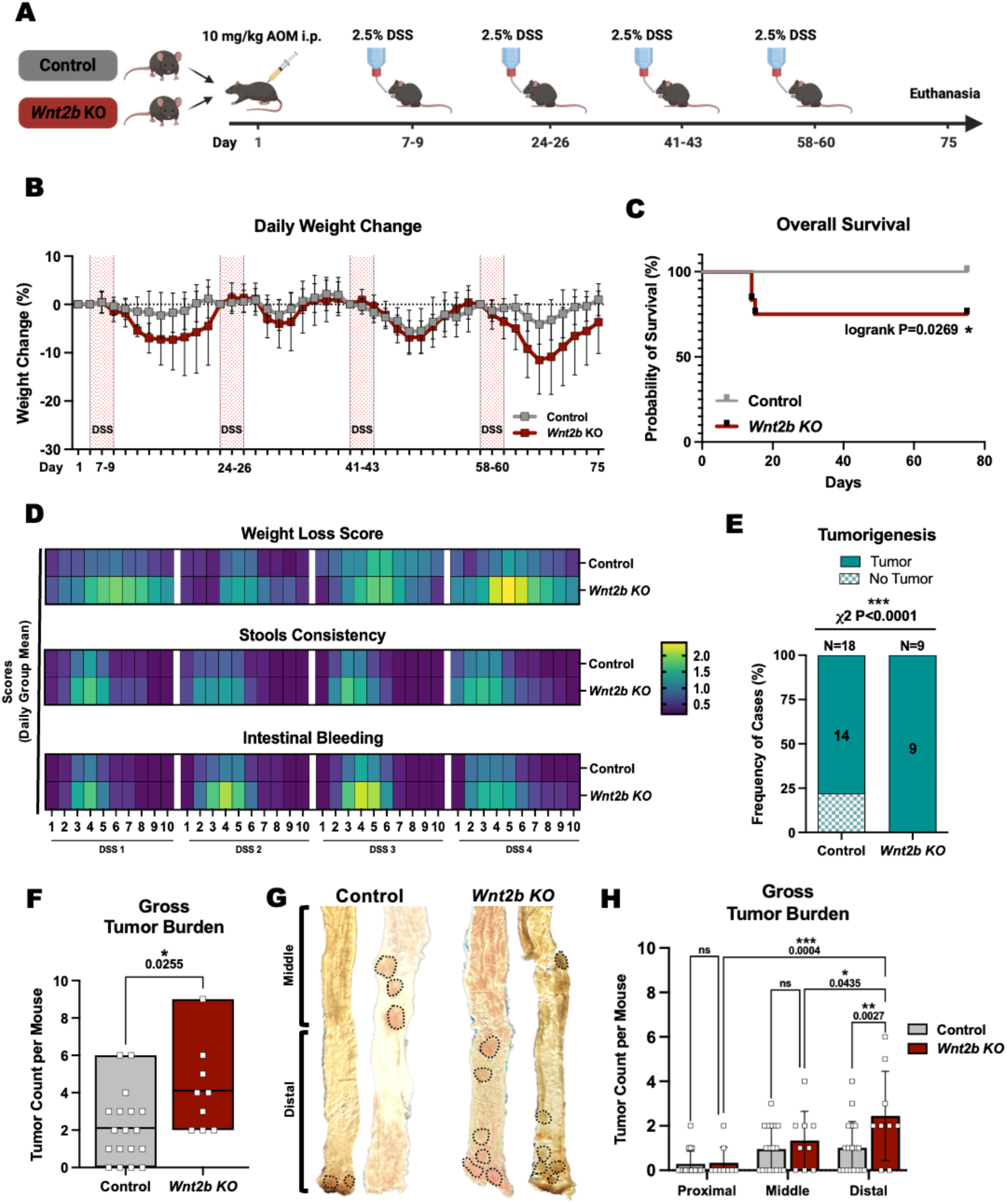
– *Wnt2b KO* Mice Are More Susceptible to AOM/DSS-induced CAC. (**A**) Adapted AOM/DSS protocol with sex and age-matched control (n=18) and *Wnt2b KO* (n=12) mice receiving 10mg/kg of Azoxymethane (AOM) at day 1, followed by four cycles of 2.5% Dextran Sodium Sulfate (DSS) in the drinking water for 3 days, with a two-week recovery period between each cycle. (**B**) Daily group mean ± SD body weight change. (**C**) Kaplan–Meier survival analysis of control (n=18) and *Wnt2b KO* (n=12) mice following AOM/DSS treatment using a log-rank test (P=0.0269). (**D**) Heat map showing the daily mean of weight loss, stool consistency, and intestinal bleeding for the first 10 days of each DSS cycle. (**E**) Frequency of animals displaying tumors during gross assessment in control (n=18) and *Wnt2b KO* (n=9) groups using chi-square test (P<0.0001). (**F**) Box plots expressing group means ± SDs of tumor burden per mouse comparing control and *Wnt2b KO* mice with a two-tailed Mann-Whitney U test (P=0.0255). (**G**) Representative macroscopic view of the intestines of the control and *Wnt2b KO* mice highlights the tumors throughout the colon. (**H**) Graph bars expressing mean ± SD of regional tumor burden per mouse in different areas of the colon, with a Two-Way ANOVA comparing control and *Wnt2b KO* (P-value shown).

### *Wnt2b KO* Mice Have Enhanced Inflammation-Driven Tumorigenesis

Next, we performed a histopathological assessment to evaluate the integrity of the colonic tissue after AOM/DSS treatment. *Wnt2b KO* mice presented slightly higher overall histopathological scores (**Fig. 2A**) due to considerably more tissue damage than controls (**Fig. S2A-F**). Still, both groups were comparable when examining inflammation and epithelial damage scores separately (**Fig. S2G-H**). On the other hand, the two *Wnt2b KO* mice that died after the first DSS cycle displayed severe intestinal damage, with the mice losing almost the whole epithelial lining and displaying increased immune cell infiltration (**Fig. 2B**). This finding corroborates our previous work and highlights that WNT2B LOF reduces tissue resilience in early colitis injury (8). Moreover, the histological assessment confirmed that *Wnt2b KO* animals that survived the entire protocol displayed a significantly greater tumor burden than control animals (**Fig. 2C, D**). Males are expected to develop more tumors than female mice undergoing AOM/DSS treatment (24). Indeed, the males exhibited a more significant tumor burden in both groups, but *Wnt2b KO* males developed significantly more tumors than control littermates (**Fig. 2E**). Furthermore, only 67% (8/12) control females developed one or two tumors on average, while all the *Wnt2b KO* females (5/5) developed three or more tumors, suggesting that WNT2B LOF may affect or supersede the estrogen receptor signaling that has been shown to protect the females in the AOM/DSS model (25). Moreover, the total tumor area was significantly larger in *Wnt2b KO* mice than in control littermates (**Fig. 2F**), which was directly associated with the tumor multiplicity (**Fig. 2G**). We also verified expression levels of three key regulators of tumor-promoting inflammation, TNFα, IL-1β, and IL-6. As expected, all three targets were more highly expressed in AOM/DSS-treated mice than in untreated animals (**Fig. 2H-J**). However, the levels of IL-6 expression were also significantly higher in the *Wnt2b* KO mice compared to the controls after AOM/DSS (**Fig. 2J**).

**Fig. 2.**
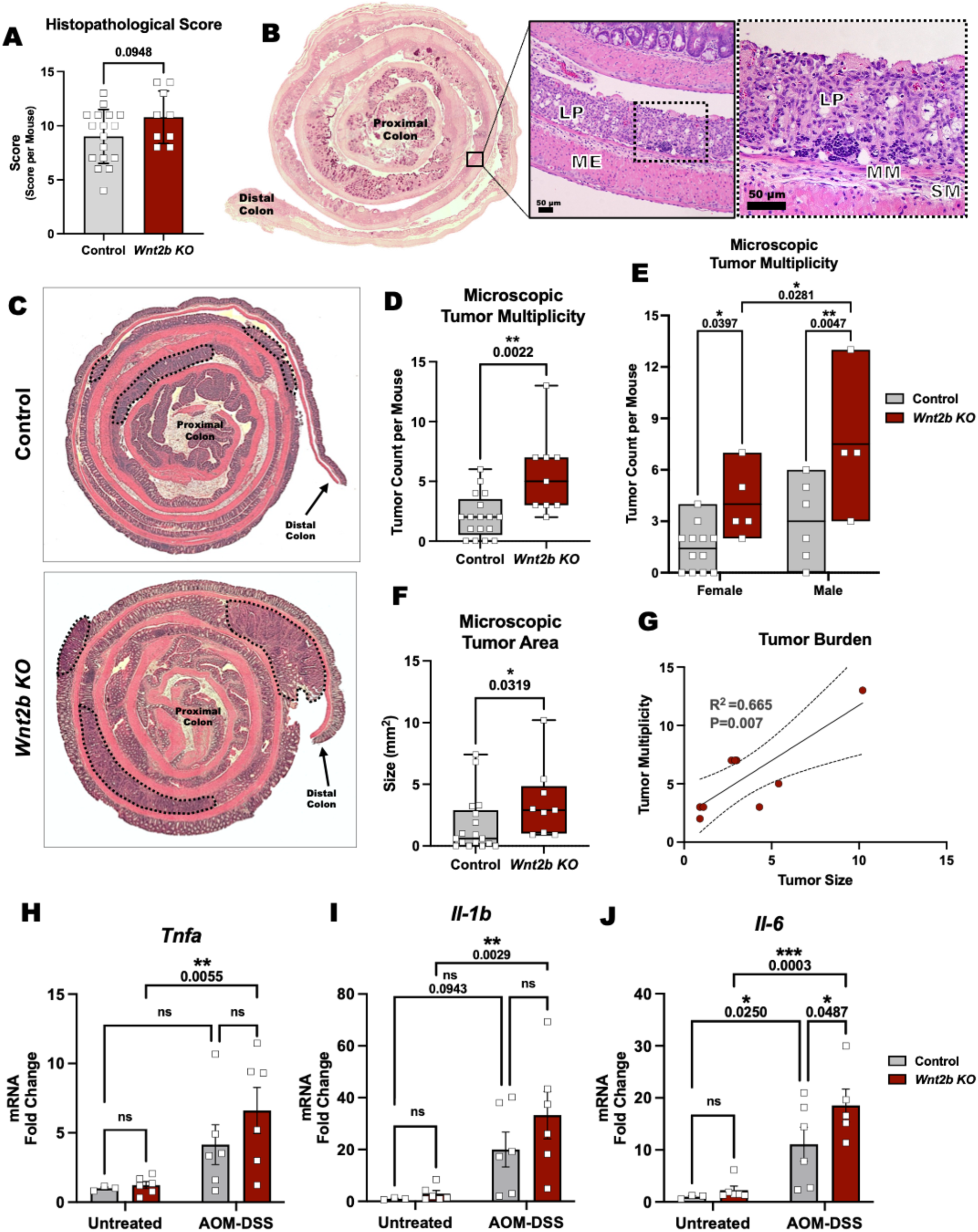
– *Wnt2b KO* Mice Have Enhanced Inflammation-Driven Tumorigenesis. (**A**) Overall histopathological scores from control and *Wnt2b KO* mice compared by a Two-tailed Student t-test (P=0.0948). (**B**) Representative PAS-stained Swiss-roll showing severe goblet cell depletion in *Wnt2b KO* mice that died after the first DSS cycle. Insets from a replicate H&E-stained slide highlighting epithelial lining loss and immune cell infiltration. Scale bars 50 µm. (**C**) Representative H&E-stained Swiss-rolls from control (top panel) and *Wnt2b KO* mice (bottom panel), with dotted lines highlighting tumor areas. (**D**) Box plots and whispers expressing tumor count per mouse in control (n=18) and *Wnt2b KO* (n=9) groups compared by a two-tailed Mann-Whitney U test (P=0.0022). (**E**) Floating bars expressing mean ± SDs comparing the sex-related differences in tumor counts via Two-way ANOVA (P-value shown). (**F**) Box plots and whispers expressing average tumor area (mm^2^) per mouse in control (n=18) and *Wnt2b KO* (n=9) groups compared by a two-tailed Mann-Whitney U test (P=0.0319). (**G**) Linear regression analysis of correlation between tumor size and tumor multiplicity (R^2^=0.665; P=0.0007). (**H-J**) Graph bars expressing mean ± SDs of *Tnfa, Il-1b,* and *Il-6* mRNA expression levels in untreated and AOM/DSS-treated samples from control (n=6) and *Wnt2b KO* (n=6) mice assessed by quantitative real-time (qRT-PCR) and compared by a Two-way ANOVA (P-value shown). Abbreviations: LP – lamina propria; MM – muscularis mucosae; SM – submucosa; ME – muscularis externa.

### *Wnt2b KO* Mice Have Increased Adenoma-Carcinoma Progression

Neoplastic areas are identified as flat plaque-like epithelial lesions presenting abnormal growth and pronounced nuclear alteration. Low-grade dysplasia adenomas (LGD) were determined by simple glandular architecture, with elongated and crowded crypts presenting hyperchromatic nuclei but maintaining polarity with the basement membrane (**Fig. 3A**). High-grade dysplasia adenomas (HGD) were characterized by glands displaying cribriform appearance with nuclear atypia and loss of nuclear polarity (**Fig. 3B**). In contrast, intraepithelial adenocarcinomas (CA) presented dense glandular lesions with tubular and/or villous structures, reduced goblet cells, nuclear atypia, with desmoplastic stroma and dirty necrosis (**Fig. 3C**). Although *Wnt2b KO* mice exhibited a larger total tumor burden due to higher number of small tumors (**Fig.S2I)**, the frequency of tumors with 1-2 mm^2^ was comparable between the two groups when comparing individual tumors (**Fig. 3D**). On the other hand, *Wnt2b KO* mice had a significantly higher frequency of HGD adenomas, and an absence of lesions showing no dysplasia (ND), compared to control littermates (**Fig. 3E**), resulting in an enhanced adenoma-carcinoma progression in *Wnt2b KO* mice (**Fig. 3F**). We next investigated whether increased tumorigenesis in *Wnt2b KO* mice would cause a compensatory hyperactivation of the canonical WNT pathway. Surprisingly, the expression of the β-catenin-target genes *Lef1, Axin2, Lgr5,* and *Myc* was comparable between *Wnt2b KO* mice and control littermates in the distal colon samples from the AOM/DSS-treated animals (**Fig. 3G**).

**Fig. 3.**
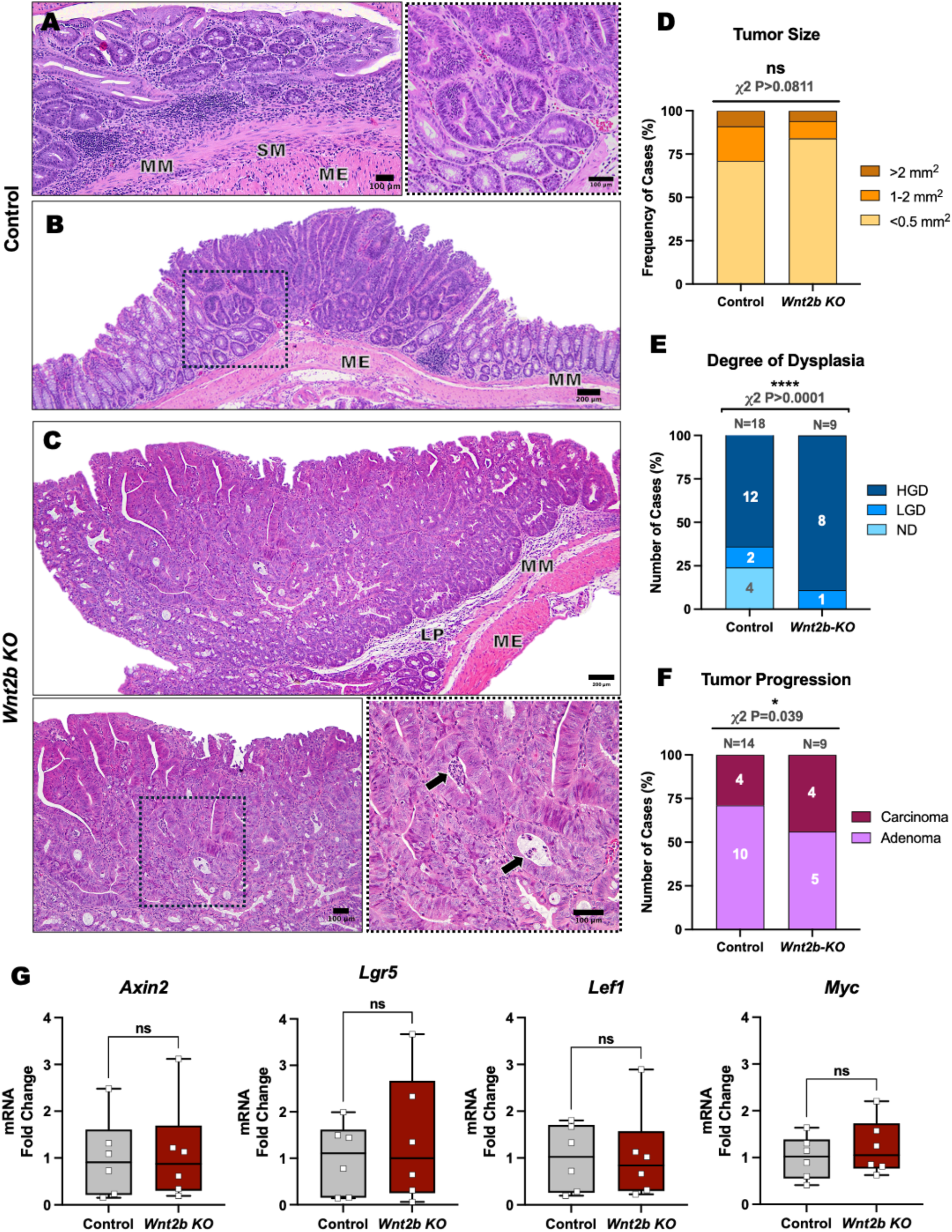
– *Wnt2b KO* Mice Have Increased Adenoma-Carcinoma Progression. (**A-C**) Representative H&E-stained sections. (**A**) Low-grade and (**B**) high-grade dysplastic adenomas from control mice, with a dashed box highlighting dysplastic glandular structures. (**C**) Intraepithelial adenocarcinoma from *Wnt2b KO* mice, with inset highlighting desmoplastic stroma and dirty necrosis (arrow). H&E staining was used to assess (**D**) tumor size, (**E**) degree of dysplasia, and (**F**) tumor staging in control and *Wnt2b KO* mice, which were compared using Fisher’s exact test (P-value shown). (**G**) Graph bars expressing mean ± SDs of *Axin2, Lgr5, Lef1,* and *Myc* mRNA expression levels in AOM/DSS-treated control (n=6) and *Wnt2b KO* (n=6) mice assessed by quantitative real-time (qRT-PCR) and compared by a Two-tailed Student t-test (P-value non-significant). Abbreviations: LP – lamina propria; MM – muscularis mucosae; SM – submucosa; ME – muscularis externa; HGD – high-grade dysplasia; LGD – low-grade dysplasia; ND – no dysplasia.

### WNT2B LOF Increases Susceptibility to Sporadic CRC Induced by AOM

Because our CAC mouse model induced carcinogenesis through a combination of mutagenesis (AOM) and inflammation (DSS), we next sought to evaluate how WNT2B LOF affects colorectal tumorigenesis without inducing an altered inflammatory response. Thus, we challenged the mice with six weekly injections of AOM without DSS (**Fig. 4A**). Unlike the AOM/DSS model, we did not observe a significant difference in weight loss comparing *Wnt2b KO* mice and control littermates (**Fig. 4B**). Interestingly, females lost more weight than males, which was more substantial among control animals (**Fig. S3A-B**). Still, no difference was observed when comparing control and *Wnt2b KO* animals by sex (**Fig. S3C-D**). Despite comparable weight loss, the survival was significantly decreased for *Wnt2b KO* mice compared to controls (**Fig. 4C**). Strikingly, 63% (5/8) of *Wnt2b KO* animals presented severe rectal prolapse (**Fig. 4D and Fig. S3E**). In contrast, control animals did not show any signs of a systemic disease (**Fig. 4D**). Interestingly, female *Wnt2b KO* mice were significantly more affected by rectal prolapse than males in this model (**Fig. 4E**). Our macroscopic assessment revealed that all five *Wnt2b KO* animals (100%) that survived the 18-week experimental protocol developed three or more tumors. In contrast, only three out of twelve (25%) control mice developed at least three tumors (**Fig. 4F**, with representative histology in **Fig. 4G**). The tumors were predominantly observed in the distal colon (**Fig. 4H-I**). No significant differences were observed for colon length or spleen weight (**Fig. S3F-G**). These findings indicate that WNT2B LOF predisposes mice to tumorigenesis even in the absence of external inflammation with DSS.

**Fig. 4.**
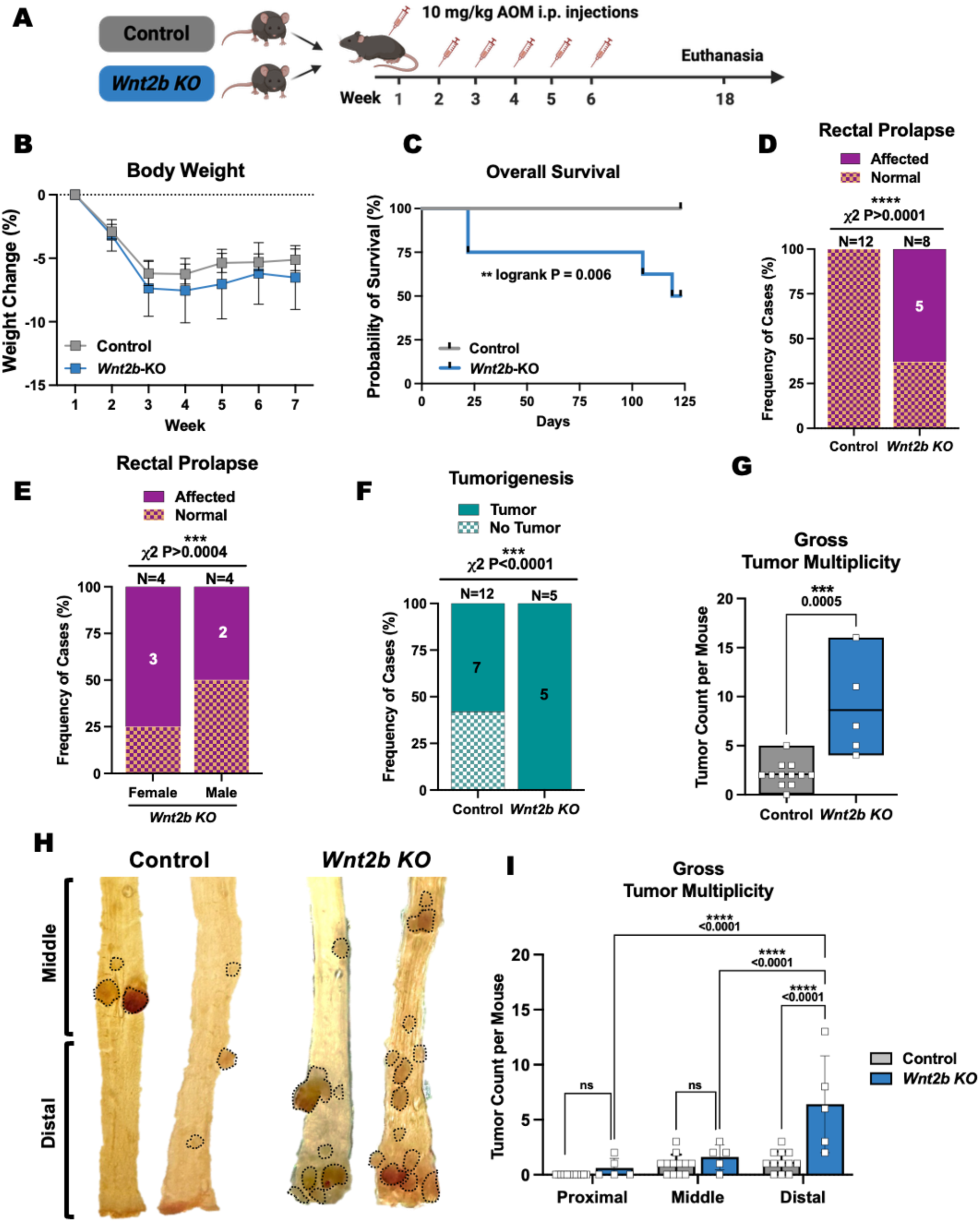
– WNT2B LOF Increases Susceptibility to Sporadic CRC. (**A**) Sporadic CRC model with sex and age-matched control (n=12) and *Wnt2b KO* (n=8) mice receiving six weekly intraperitoneal injections of 10mg/kg AOM, followed by twelve weeks for tumor development under standard conditions. (**B**) Graph expressing weekly group mean ± SDs body weight variation, with no significant difference between groups. (**C**) Kaplan– Meier survival analysis of control (n=12) and *Wnt2b KO* (n=8) mice following AOM injections using a log-rank test (P=0.006). Frequency of cases of rectal prolapse compared by group (**D**) or by sex (**E**) using Fisher’s exact test (P-value shown). (**F**) Frequency of animals displaying tumors during gross assessment in control (n=12) and *Wnt2b KO* (n=5) mice that survived until the end of the experiment was compared by a Fisher’s exact test (P<0.0001). (**G**) Floating bars expressing group means ± SDs of tumor multiplicity per colon in control (n=12) and *Wnt2b KO* (n=5), calculated by a Two-tailed Mann-Whitney U test (P=0.0005). (**H**) A representative macroscopic view of the distal colon of the control and *Wnt2b KO* mice highlights the tumors (dotted lines). (**I**) Graph bars expressing mean ± SD of regional tumor multiplicity per mouse in different areas of the colon, with a Two-Way ANOVA comparing control and *Wnt2b KO* (P-value shown).

### *Wnt2b KO* Mice Have Greater Tumor Area in the Sporadic CRC Model

Because tumor burden can reflect both tumor numbers and tumor size, we next analyzed the total tumor area of each mouse. The histopathological analysis revealed that *Wnt2b KO* mice have enhanced colonic injury after AOM treatment. Interestingly, we observed slightly higher inflammation scores for *Wnt2b KO* mice (**Fig. S2G**). Still, it did not reach statistical significance when compared to control littermates (**Fig. 5A**). However, epithelial damage scores were significantly higher in *Wnt2b KO* mice due to larger areas of hyperplasia and dysplasia (**Fig. 5B**). Consequently, overall histopathological scores were significantly higher in *Wnt2b KO* mice compared to controls (**Fig. 5C**). *Wnt2b KO* mice had comparable adenomas to controls. In contrast, carcinomatous areas were larger compared to control littermates (**Fig. 5D-F**). Strikingly, the total neoplastic coverage areas were above 5 mm^2^ per colon in some *Wnt2b KO* animals (**Fig. 5G**), which was positively associated with the number of tumors (**Fig. 5G**).

**Fig. 5.**
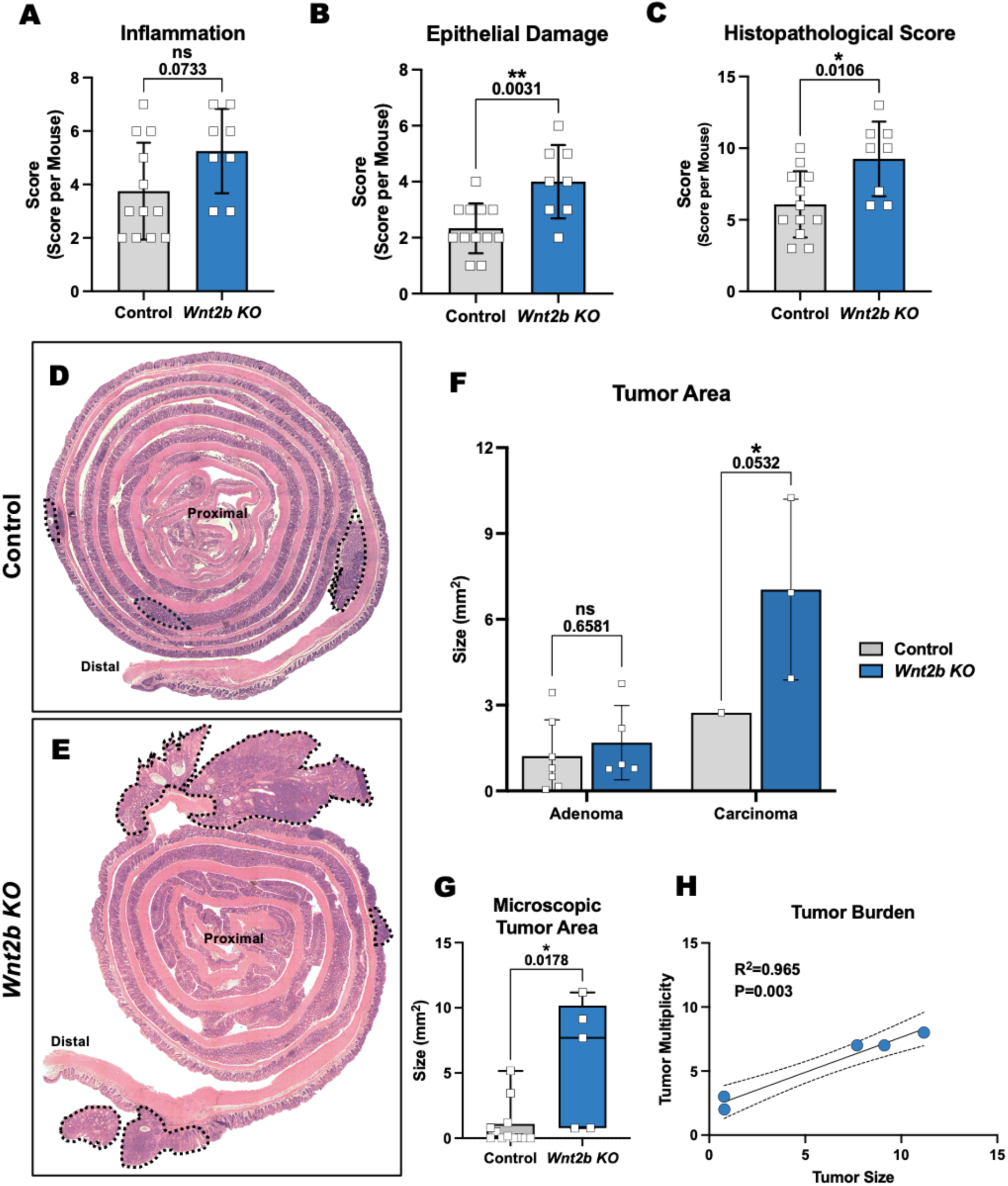
– *Wnt2b KO* Mice Have Greater Tumor Area in Sporadic CRC Model. Histopathological assessment was performed using H&E staining and graph bars expressing mean ± SD of (**A**) inflammation (immune infiltration), (**B**) epithelial damage (ulceration and hyperplasia), and (**C**) overall histopathological scores from control (n=12) and *Wnt2b KO* (n=8) mice compared by a Two-tailed Student t-test (P-value shown). Representative H&E-stained Swiss-rolls from (**D**) control and (**E**) *Wnt2b KO* mice, with dotted lines highlighting tumor areas. (**F**) Graph bars expressing mean ± SD of total adenoma and adenocarcinoma areas in control (n=12) and *Wnt2b KO* (n=5) mice compared by a one-way ANOVA (P-value shown). (**G**) Box plots and whispers expressing average tumor area (mm^2^) per mouse in control (n=12) and *Wnt2b KO* (n=5) groups compared by a Two-tailed Mann-Whitney U test (P=0.0178). (**H**) Linear regression analysis of correlation between tumor size and tumor multiplicity (R^2^=0.965; P=0.003).

### WNT2B LOF Predisposes to the Development of Sporadic Adenocarcinoma

Next, we evaluated tumor morphology of spontaneously occurring tumors in the AOM-only model. When comparing only adenocarcinomas, control animals displayed a more homogeneous and dense cribriform glandular morphology containing mucous and neutrophils (**Fig. 6A**). In contrast, *Wnt2b KO* animals presented large adenocarcinomas (**Fig. 5F**) with heterogenous morphology, highlighted by a desmoplastic stroma with increased areas containing dirty necrosis (**Fig. 6B**). No difference was observed for class of tumor size (**Fig. 6C**), but *Wnt2b KO* mice revealed a greater frequency of lesions of high-grade dysplasia compared to controls, and an absence of no dysplasia lesions (**Fig. 6D**). Consequently, the progression from adenomas to carcinomas was significantly higher in *Wnt2b KO* animals (**Fig. 6E**), indicating that WNT2B LOF increases the susceptibility to tumor development even in the absence of induced inflammation.

**Fig. 6.**
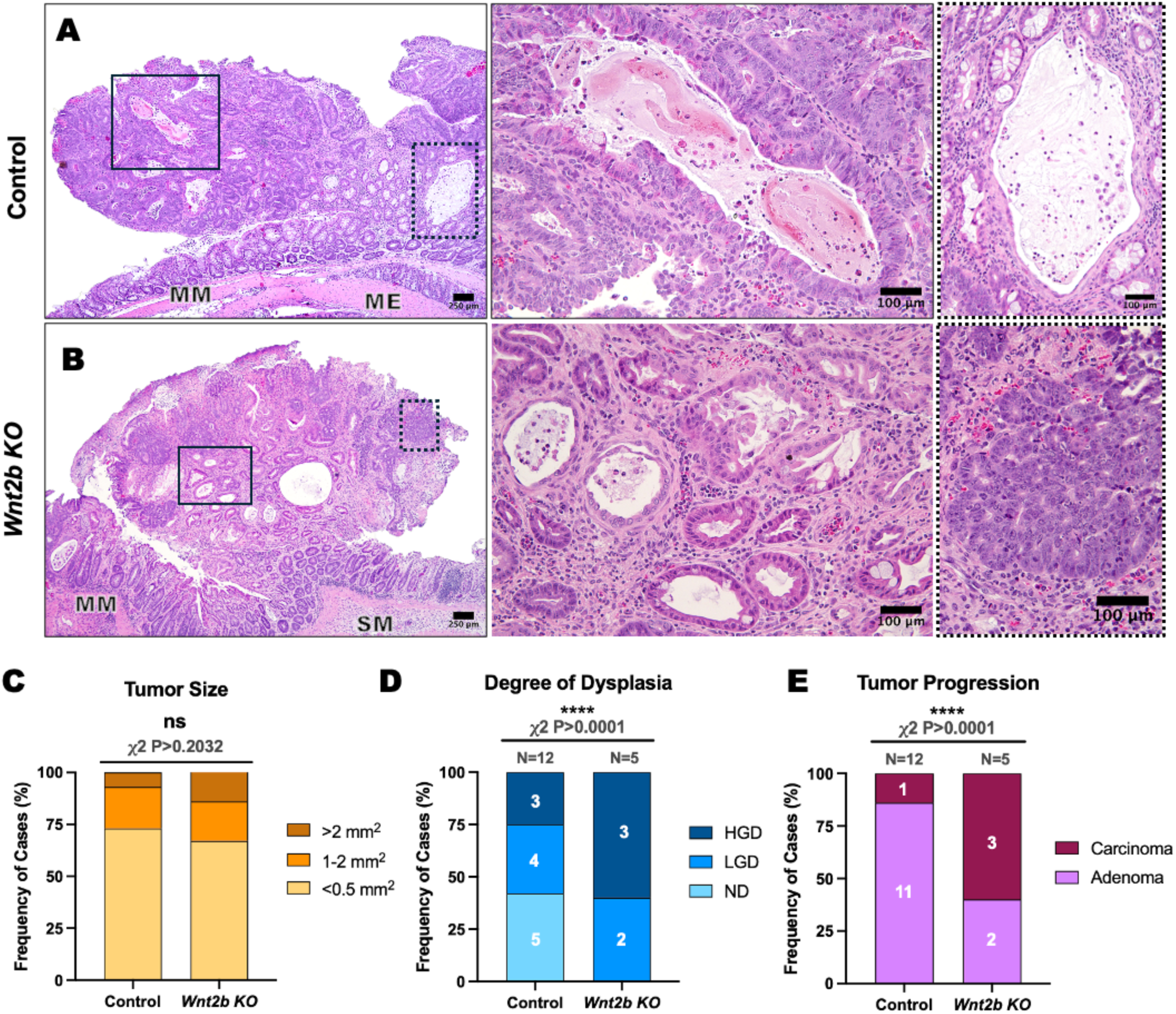
– WNT2B LOF Predisposes to the Development of Sporadic Adenocarcinoma. Representative H&E-stained intraepithelial adenocarcinoma in (**A**) control and (**B**) *Wnt2b KO* mice. Scale bars 250 µm. Enlarged images highlight the differences in secretory and immune cell content inside glandular structures, which display heterogeneous epithelial and stromal morphologies. Scale bars 100 µm. H&E staining was used to assess (**C**) tumor size, (**D**) degree of dysplasia, and (**E**) disease staging. Control (n=12) and *Wnt2b KO* (n=5) groups were compared using Fisher’s exact test (P-value shown). Abbreviations: MM – muscularis mucosae; SM – submucosa; ME – muscularis externa; ND – no dysplasia; LGD – low-grade dysplasia; HGD – high-grade dysplasia.

### WNT2B Loss Predicts a Worse Prognosis for Patients with CRC

Next, we used the TCGA dataset of human sporadic colon cancer (COAD) to understand how *WNT2B* expression relates to human CRC using GEPIA and TNMplot web-based platforms. We found that *WNT2B* expression was reduced in colon cancer patients compared to healthy control samples (**Fig. 7A**). However, no significant difference in *WNT2B* levels was observed across the four main stages of the disease (**Fig. 7B**), suggesting that reduced levels of *WNT2B* expression is a continuous event during tumor progression or at least after tumor initiation. Furthermore, we found that low levels of *WNT2B* were associated with reduced relapse-free survival (**Fig. 7C**), especially in early stages of the disease (**Fig. S4A-B**). We then investigated the frequency of copy number variation (CNV) in the *WNT2B* gene using cBioPortal. We found that mutations leading to homozygous (deep) and heterozygous (shallow) deletions were observed in 29% of the profiled samples combined (**Fig. 7D**). Of note, these mutations lead to *WNT2B* LOF and were significantly associated with reduced overall survival compared to patients carrying the consensus sequence (**Fig. 7E**). By contrast, mutations leading to homozygous or heterozygous amplifications, all culminating in GOF, were highly frequent in *WNT2* (42%), *WNT3* (19%), *WNT5A* (11%) (**Fig. 7F**). We also observed that *WNT2B* gene is hypermethylated in CRC samples (**Fig. S4C**), which is negatively associated with gene expression (**Fig. S4D**). Together, these findings suggest that WNT2B expression is lost during early stages of tumor development and is associated with worse patient prognosis. This supports our hypothesis that WNT2B has a protective role against the development of CRC.

**Fig. 7.**
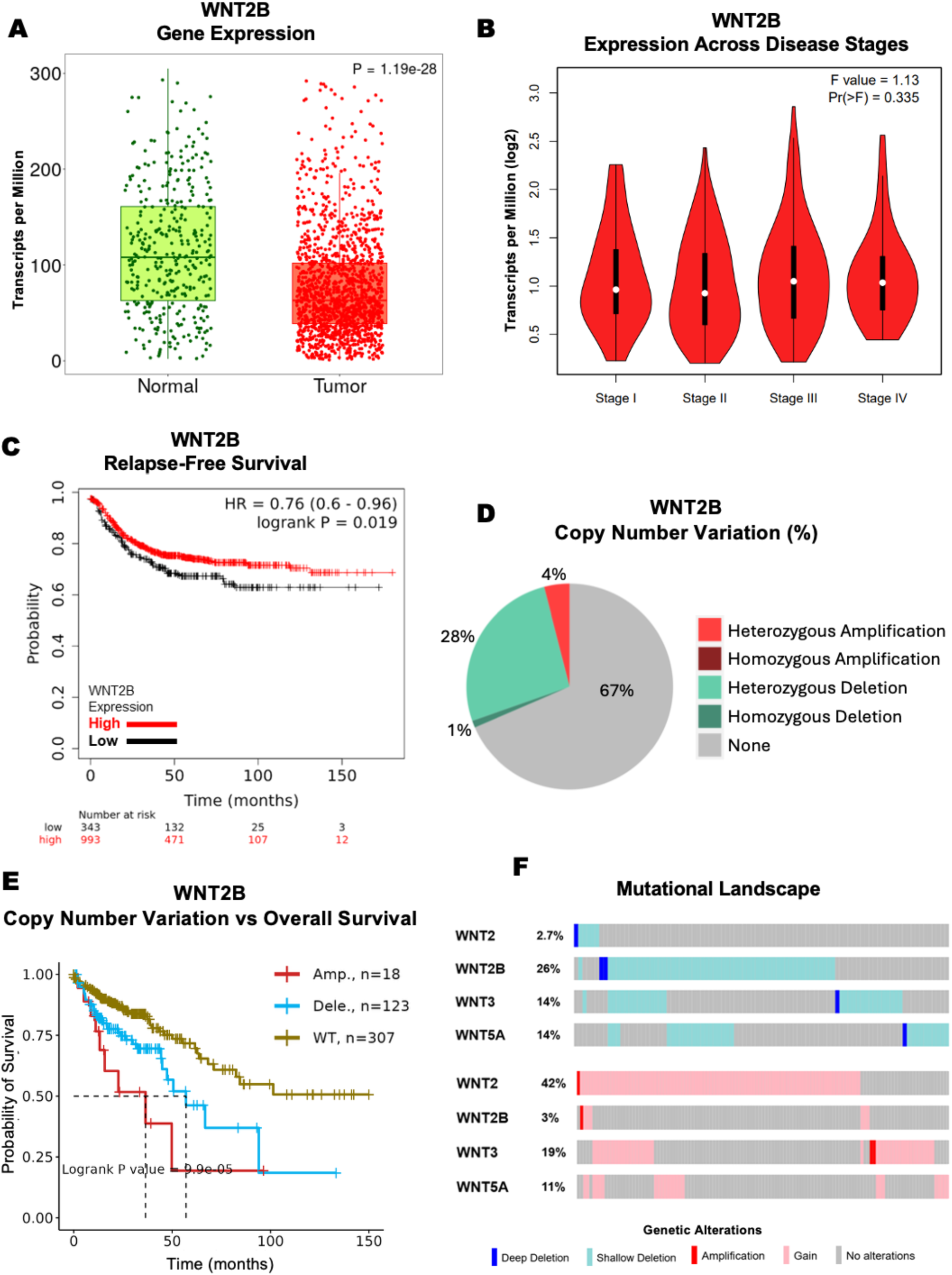
– WNT2B Loss Predicts a Worse Prognosis for Patients with CRC. (**A**) WNT2B expression levels in normal (n=377) and colon cancer samples (n=1450) from TCGA’s colon adenocarcinoma (COAD) dataset. (**B**) WNT2B expression levels in colon cancer samples were compared across the four disease stages (Pr(>F) = 0.335). (**C**) Kaplan–Meier curves show the association between high (red) and low (black) WNT2B expression levels and relapse-free survival of patients from all stages calculated by a Mantel-Cox test (P=0.019) (**D**). Pie plot showing the constitution of copy number variation in the WNT2B gene profiled in the CRC samples (COAD) dataset. (**E**) Kaplan– Meier curves show the association between WNT2B mutation and patient survival calculated by a Mantel-Cox test (P=0.00099). (**F**) Frequency of copy number alteration in WNT2, WNT2B, WNT3, and WNT5A gene profiles in CRC samples from the COAD dataset.

## Discussion

Our study demonstrated that WNT2B LOF increases susceptibility to sporadic and colitis-associated cancers. *Wnt2b KO* mice exhibited a more significant tumor burden and larger areas of highly dysplastic adenomas and adenocarcinomas than control littermates, which culminated in reduced survival and a high adenoma-to-carcinoma progression in both experimental models. Moreover, we found that WNT2B expression was reduced in human CRC samples due to a high frequency of mutations and gene hypermethylation, which was significantly linked to a worse patient prognosis. While WNT signaling has been widely explored in CRC, this has mainly been in the context of b-catenin activation by canonical WNTs (5). Our findings indicate that WNT2B is protective against tumor development, and that loss of WNT2B does not affect canonical b-catenin targets, which suggests that WNT2B may have b-catenin-independent or pleiotropic functions. Indeed, while *WNT2B* expression is decreased in CRC samples, expression of other prominent intestinal WNTs (e.g., WNT3 and WNT5A) is increased (**Fig.S4**), and loss of these molecules has been shown to inhibit tumorigenesis (26, 27). Importantly, WNT3 is the prototypical β-catenin-dependent Wnt and WNT5A is the prototypical β-catenin-independent Wnt, and since they both seem to work similarly in terms of cancer susceptibility, it suggests that the opposite activity of WNT2B in this regard may be more complex than the primary signaling variety. The mechanism of action of WNT2B in protecting against CRC may instead overlap with its function in preventing colitis in the colon, possibly via modulation of the immune response. Indeed, levels of IL-6 were significantly higher in *Wnt2b KO* animals compared to control littermates. Notably, IL-6 acts as a mitogen and angiogenic factor in the microenvironment, driving tumor cell proliferation, survival, and epithelial-mesenchymal transition (28, 29). Furthermore, high serum or tissue IL-6 levels correlate with worse patient prognosis (30, 31).

CRC arises from genetic alterations and the inability of epithelial cells to recover from damage caused by different stressors, such as infection and chronic inflammation (32). AOM promotes mutagenesis through metabolic activation and DNA damage, but it also synergizes with inflammation, which enhances mutagenesis by increasing reactive oxygen and nitrogen species (12, 14, 33). Since *Wnt2b KO* mice were previously shown to have enhanced susceptibility to acute inflammation due to increased baseline inflammation, an enhanced inflammatory response after AOM treatment may explain why the tumor burden in *Wnt2b KO* mice was even more remarkable than that observed in control littermates in our sporadic CRC model. In addition, since WNT2B deletion was associated with increased cancers and did not impact canonical WNT target genes, targeting WNT2B may hold potential as a therapeutic intervention in IBD and CRC.

While our study provides compelling evidence that WNT2B loss exacerbates CRC development, we acknowledge that our AOM/DSS and AOM-only mouse models may not fully recapitulate the complexity of human CRC, particularly sporadic cases. Moreover, using a global knockout model makes it difficult to isolate the specific role of WNT2B within the intestinal epithelium or tumor microenvironment and determine the precise mechanism by which WNT2B protects the colon against tumorigenesis. While suggestive, the human data derived from TCGA are correlative and not supported by functional validation or independent patient cohorts. Still, these findings support our hypothesis that WNT2B has a protective role against CRC development, and future studies will need to address these limitations.

In sum, we demonstrate the relevance of WNT2B in sporadic and colitis-associated CRC using genetically modified mice in vivo. Our study confirms that WNT2B is protective against dysplasia and CRC development, both in response to mutagenesis and chronic inflammatory stress. Moreover, human data from CRC databases paralleled the mouse model findings, suggesting that WNT2B may be important for resistance to human CRC.

## Acknowledgments

We thank Chidera Emeonye and Nicholas Makogonov for their technical assistance with the mouse colony and tissue recovery.

## Author contributions

L.F.S.O. conceptualized the project, performed the experiments, and wrote and reviewed the manuscript. Y.W. managed the mouse colony and contributed to mouse and qPCR experiments. S.R., J.K., and C.A. contributed to mouse experiments. V.S.D. contributed to the microscopy analysis. J.P. performed assessments and contributed critically to the histopathological aspects of the work. J.O.M. and D.L.C. contributed to formal analysis and experimental design. D.T.B. contributed to formal analysis, experimental design, and critically reviewed and edited the manuscript. A.E.O. conceptualized and supervised the project, contributed to formal analysis, and critically reviewed and edited the manuscript. All authors critically reviewed the manuscript and approved the final manuscript version.

## Abbreviations

ANOVA: Analysis of variance
AOM: Azoxymethane
APC: Adenomatous polyposis coli
ARCH: Animal resources at Boston Children’s Hospital
CAC: Colitis-associated cancer
CAFs: Cancer-associated fibroblasts
COAD: Colon adenocarcinoma
CMV: Cytomegalovirus
CNV: Copy number variation
CRC: Colorectal cancer
DAI: Disease activity index
DKK: Dickkopf
DSS: Dextran sodium sulfate
FAP: Fibroblast activation protein
FFPE: Formalin-fixed, paraffin-embedded
GOF: Gain-of-function
H&E: Hematoxylin and eosin
HET: Heterozygous
HGD: High-grade dysplasia
HOM: Homozygous
IACUC: Institutional animal care and use committee
IBD: Inflammatory bowel disease
IL-1: Interleukin 1
IL-6: Interleukin 6
ISC: Intestinal stem cells
LEF1: Lymphoid enhancer-binding factor 1
LGD: Low-grade dysplasia
LGR5: Leucine-rich repeat-containing G-protein coupled receptor 5
LOF: Loss-of-function
MYC: MYC proto-oncogene
MDSCs: Myeloid-derived suppressor cells
NOTUM: Palmitoleoyl-protein carboxylesterase
PAS: Periodic acid-Schiff
PCR: Polymerase chain reaction
SD: Standard deviation
TCGA: The Human Genome Atlas
TNFα: Tumor necrosis factor-alpha
TME: Tumor microenvironment
WIFI1: WNT inhibitory factor 1
WNT: Wingless-related integration site protein
WT: Wild-type

---

**Supplementary Table 1.** Key Resources Table.

| REAGENT | SOURCE | IDENTIFIER |
| --- | --- | --- |
| ***Chemicals*** |  |  |
| Azoxymethane (AOM) | Sigma-Aldrich | A5486 |
| Dextran Sodium Sulfate (SDS) | MP Biomedicals | 160110 |
| 10% Neutral Buffered Formalin Solution | Sigma-Aldrich | HT501128 |
| 1X Phosphate Buffered Solution, pH 7.4 | Gibco | 10010023 |
| TRIzol™ Reagent | Invitrogen | 15596018 |
| 28G Insulin Syringe 1ml | BD | 329424 |
| Ethyl Alcohol 200 Proof | Pharmco-Aaper | 111000200CSGL |
| ***Critical Commercial Assays*** |  |  |
| RNeasy Mini Kit | QIAGEN | 74104 |
| High-Capacity cDNA Reverse Transcription Kit | ThermoFisher | 4368813 |
| TaqMan™ Universal PCR Master Mix | ThermoFisher | 4364338 |
| ***Oligonucleotides*** |  |  |
| 18S | ThermoFisher | Mm03928990_g1 |
| Wnt2b | ThermoFisher | Mm00437330_m1 |
| Lef1 | ThermoFisher | Mm00550265_m1 |
| Axin2 | ThermoFisher | Mm01265779_m1 |
| Lgr5 | ThermoFisher | Mm00438890_m1 |
| Myc | ThermoFisher | Mm00487804_m1 |
| IL-6 | ThermoFisher | Mm00446190_m1 |
| TNFa | ThermoFisher | Mm00443258_m1 |
| IL-1B | ThermoFisher | Mm00434228_m1 |
| ***Software and Algorithms*** |  |  |
| GraphPad Prism 10.2.2 | GraphPad Prism | N/A |
| QuPath v0.6.0 | QuPath | N/A |

**Supplementary Table 2.** Disease Activity Index (DAI)

| Score<sup>a</sup> | Weight loss | Feces consistency | Intestinal bleeding |
| --- | --- | --- | --- |
| 0 | None | Normal | Normal |
| 1 | 0%–5% | Soft but formed | Blood traces in stool visible |
| 2 | 5%–10% | Soft but unformed | Blood traces in stool visible |
| 3 | 10%–18% | Very soft and wet | Archorrhagia |
| 4 | >18% | Diarrhea | Rectocele |

<sup>a</sup>The sum of the three subscores results in a combined score ranging from 0 (no changes) to 12 (severe disease activity).

**Supplementary Table 3.** Histopathological Assessment.

| Score<sup>a</sup> | Inflammatory Infiltrates |  | Epithelial Changes | Ulceration |
| --- | --- | --- | --- | --- |
|  | Severity | Extent |  |  |
| 1 | Minimal | Mucosa | Minimal hyperplasia |  |
| 2 | Mild | Mucosa, sometimes submucosa | Hyperplasia ± goblet cell loss ± cryptitis ± erosions |  |
| 3 | Moderate | Mucosa and submucosa | Hyperplasia ± goblet cell loss ± cryptitis ± crypt abscesses | Ulcerations |
| 4 | Marked | Transmural | Hyperplasia ± goblet cell loss ± cryptitis ± multiple crypt abscesses | Extended ulcerations ± pseudo polyps |

<sup>a</sup>The overall histopathological score is obtained from the sum of the three subscores, ranging from 1 (no changes) to 12
(severe disease activity).

