## Supplementary Figures for "Loss of WNT2B Increases Progression from Dysplasia to Colorectal Cancer"

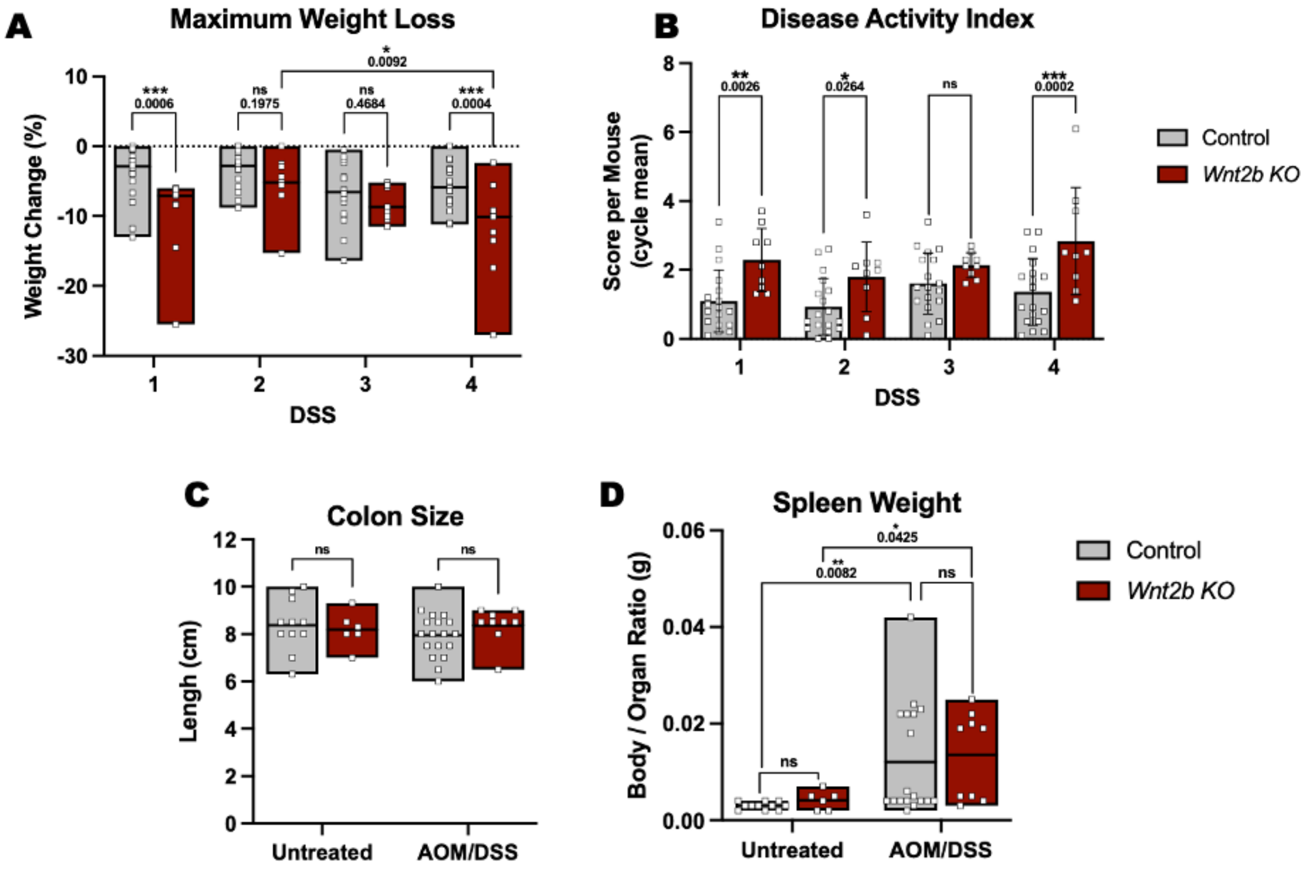


**Fig. S1 – *Wnt2b KO* Mice Are More Susceptible to AOM/DSS-induced Cancer.** (**A**) Maximum body weight loss in each DSS cycle with Two-Way ANOVA comparing control and *Wnt2b KO* (P-value shown). (**B**) Graph bars expressing mean ± SD of DAI in each DSS cycle, with a Two-Way ANOVA comparing control and *Wnt2b KO* (P-value shown). (**C**) Floating bars expressing mean ± SD colon size per mouse, with a Two-Way ANOVA comparing control and *Wnt2b KO* (P>0.05). (**D**) Floating bars expressing mean ± SD spleen weight per mouse, with a Two-Way ANOVA comparing control and *Wnt2b KO* (P-value shown

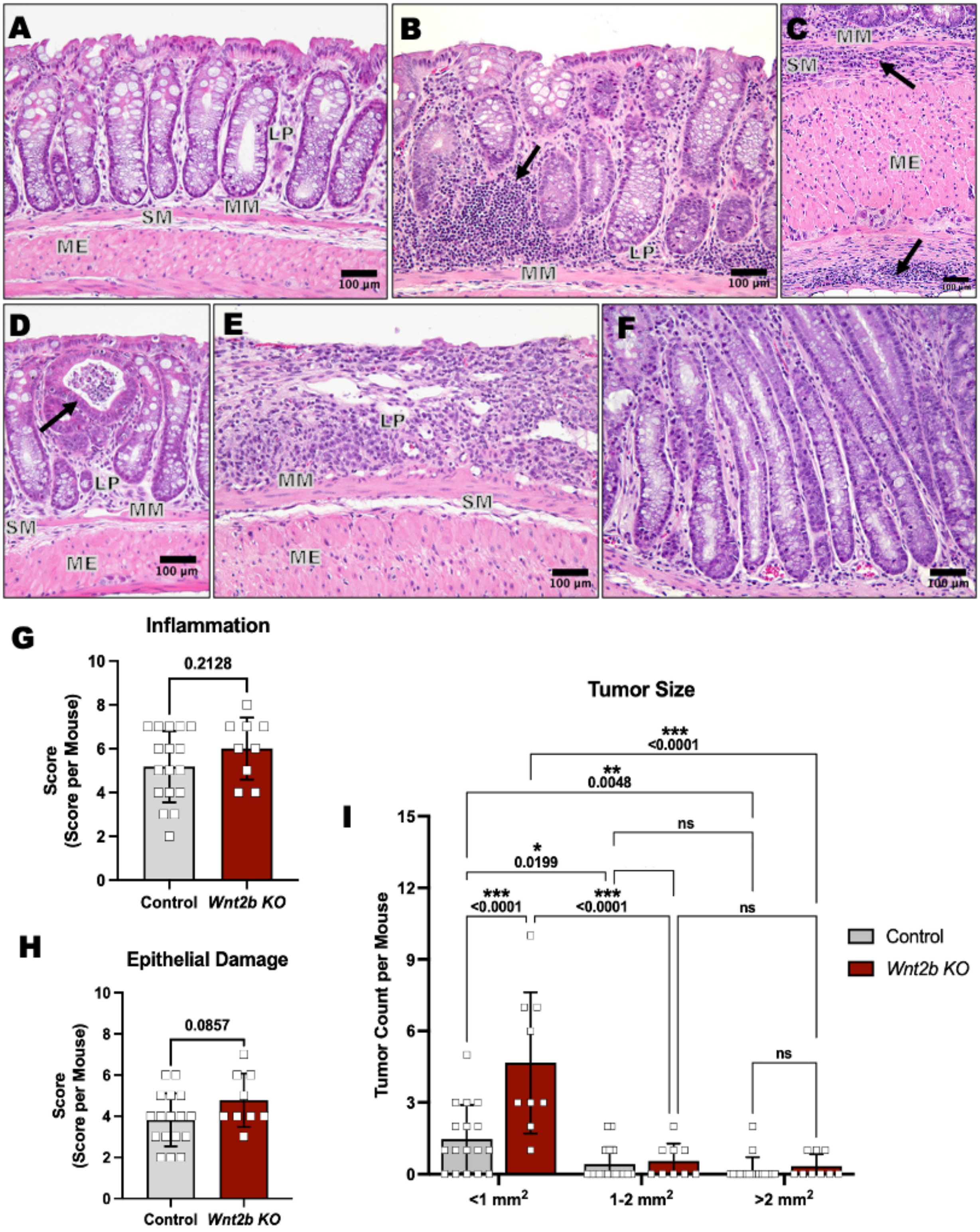


**Fig. S2 – *Wnt2b KO* Mice Have Enhanced Inflammation-Driven Tumorigenesis.** (**A-F**) Representative H&E-stained images. (**A**) Healthy colon tissue from an untreated mouse. (**B**) Mucosal immune infiltration. (**C**) Transmural immune infiltration. (**D**) Crypt abscess. (**E**) Ulceration. (**F**) Crypt hyperplasia. (**G**) Histological score of inflammation. (**H**) Histological score of epithelial damage. (**I**) Graph bars comparing tumor size in control and *Wnt2b KO* mice using Two-way ANOVA (P-value shown)

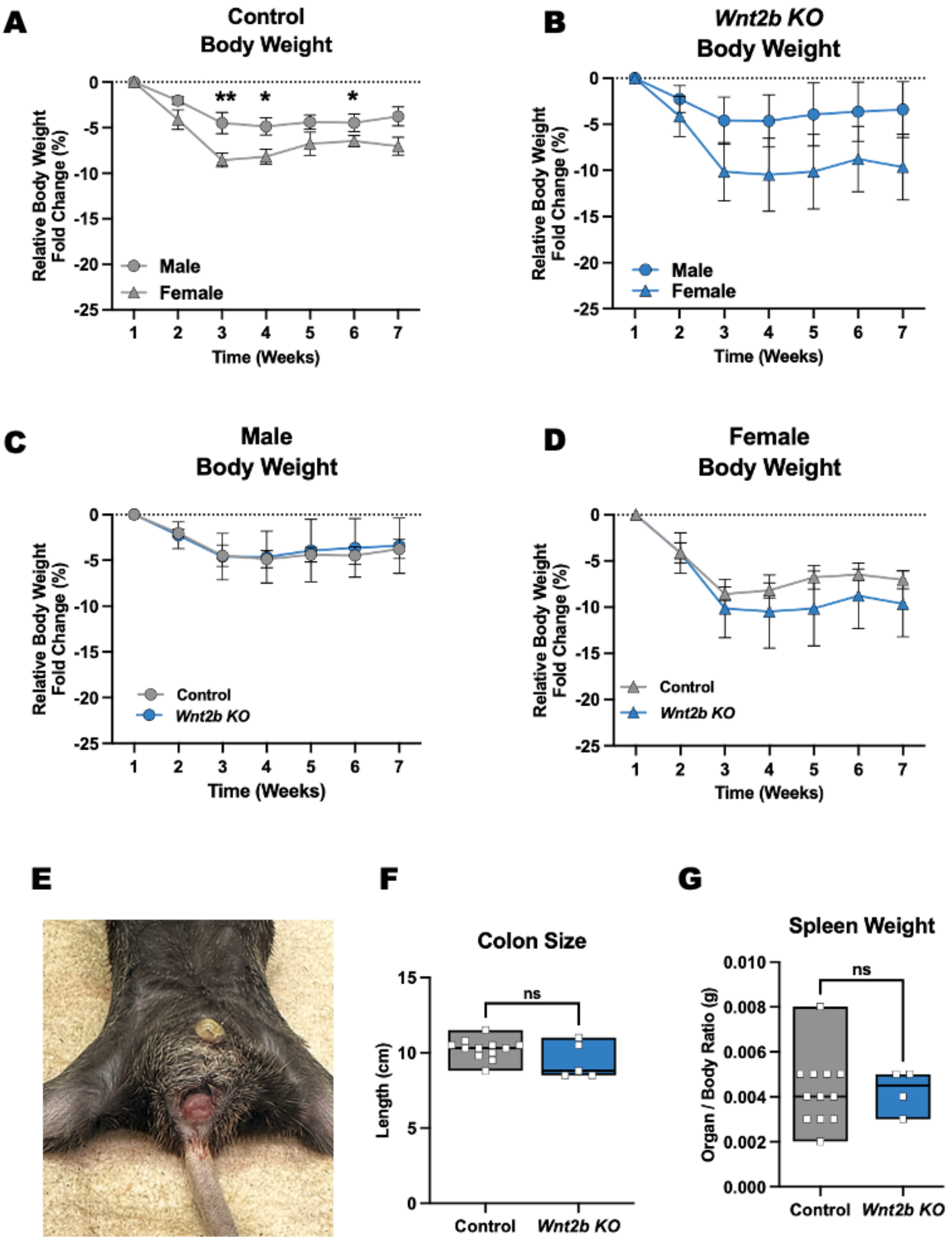


**Fig. S3 – WNT2B LOF Increases Susceptibility to Sporadic CRC.** (**A-D**) Weekly group mean ± SD body weight change. (**A**) Body weight change in control animals. (**B**) Body weight change in *Wnt2b KO* animals. (**C**) Body weight change in control and *Wnt2b KO* males. (**D**) Body weight change in control and *Wnt2b KO* females. (**E**) Rectal prolapse. (**F-G**) Floating bars expressing mean ± SD colon size and spleen weight per mouse, with a two-tailed Mann-Whitney U test comparing control and *Wnt2b KO* (P>0.05).

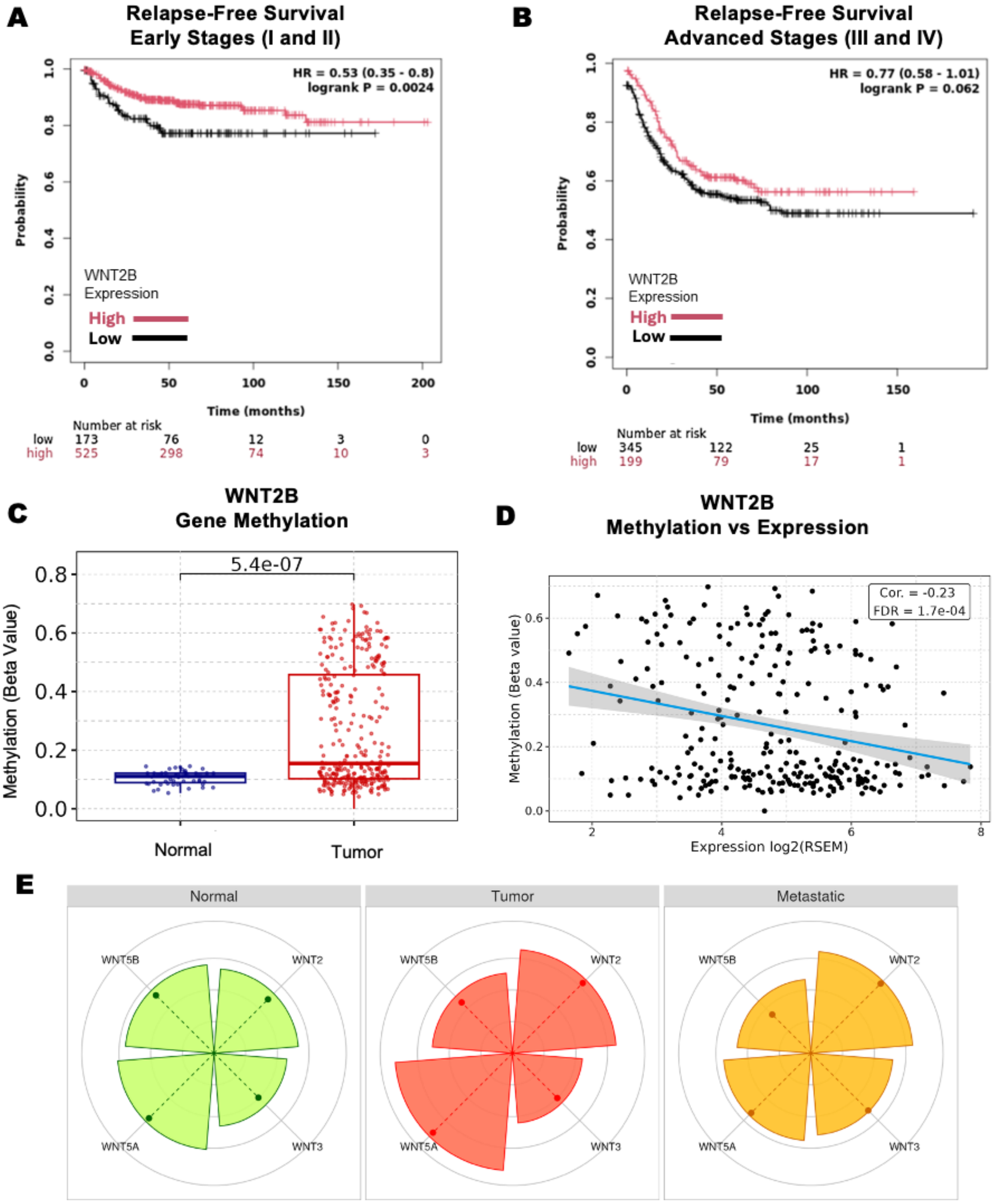


**Fig. S4 – *WNT2B* Loss Predicts a Worse Prognosis for Patients with CRC.** (**A-B**) Kaplan–Meier curves show the association between high (red) and low (black) WNT2B expression levels and relapse-free survival of patients from early stages (**A**) and late stages (**B**), calculated by a Mantel-Cox test using KM Plotter (P-value shown). (**C**) Methylation levels of the WNT2B gene and (**D**) correlation of WNT2B expression and promoter methylation produced using cBioPortal. (**E**) Expression levels of different WNT ligands in healthy, tumor, and metastatic samples using TNMplot.
